# MEK inhibitor resistance in lung cancer cells associated with addiction to sustained ERK suppression

**DOI:** 10.1101/2022.04.29.490009

**Authors:** Dylan A. Farnsworth, Yusuke Inoue, Fraser D. Johnson, Georgia de Rappard-Yuswack, Daniel Lu, Rocky Shi, Romel Somwar, Marc Ladanyi, Arun M. Unni, William W. Lockwood

## Abstract

MEK inhibitors have yielded limited efficacy in KRAS-mutant lung adenocarcinoma (LUAD) patients due to drug resistance. We established trametinib-resistant KRAS-mutant LUAD cells and describe a state of “drug addiction” in a subset of resistant cases where cells are dependent on trametinib for survival. Dependence on ERK2 suppression underlies this phenomenon whereby trametinib removal hyperactivates ERK and results in ER stress and apoptosis. Amplification of *KRAS^G12C^* occurs in drug-addicted cells and blocking mutant specific activity with AMG 510 rescues the lethality after trametinib withdrawal. Furthermore, increased KRAS^G12C^ expression is lethal to other KRAS mutant LUAD cells, consequential to ERK hyperactivation. Our study represents the first instance of this phenotype associated with *KRAS* amplification and demonstrates that acquired genetic changes that develop in the background of MAPK suppression can have unique consequence. We suggest that the presence of mutant *KRAS* amplification in patients may identify those that may benefit from a “drug holiday” to circumvent drug resistance. These findings demonstrate the toxic potential of hyperactive ERK signaling and highlight potential therapeutic opportunities in patients bearing *KRAS* mutations.

## Introduction

Activating mutations in *KRAS* occur in approximately 30% of lung adenocarcinoma (LUAD), the major molecularly-defined subtype of lung cancer (1, 2). Patients bearing tumors with *KRAS* mutations display shorter median survival, due in part to the lack of available targeted therapies (3). In contrast, for patients with tumors driven by alterations in *EGFR*, *MET*, *ALK* or *ROS1*, selective inhibitors have improved outcomes (4–6). The presence of *KRAS* mutations has also been associated with decreased benefit from chemotherapy (7), as well as overall poor prognosis. AMG 510 is the first KRAS mutant-specific agent to enter clinical trials in humans, and was recently granted Breakthrough Therapy designation from the FDA. AMG 510 is specific for KRAS with the G12C substitution, which is detected in approximately 30-40% of KRAS-mutant LUAD tumors (8). However, in a recent phase 1 trial of AMG 510, only 32% of LUAD patients (19/59) had a confirmed objective response and the median progression-free survival was 6.3 months (9). It is likely that the poor response of LUAD patients with KRAS^G12C^ to AMG 510 could be due to the ability of these cancers to quickly adapt to this targeted therapy or to the presence of pre-existing drug-resistant clones, as has been described in the context of other targeted therapies (10). Indeed, *in vitro* studies have shown that resistance to KRAS^G12C^-specific inhibitors may develop rapidly (11, 12). These early results suggest that there remains an urgent need for new therapeutic strategies for LUAD patients with KRAS-driven cancers.

One previously explored avenue for the treatment of KRAS-driven lung cancers is through the inhibition of downstream pathway effectors. The RAS-RAF-MEK-ERK (MAPK) signaling pathway is a key pathway activated by mutant KRAS and plays a critical role in cell proliferation, survival, and differentiation (13, 14). Analysis of IC50s for growth inhibition across multiple cell lines shows that when compared to KRAS-wild type cells, KRAS-mutant cell lines are most sensitive to MEK inhibitors compared to inhibitors of other cancer-associated pathways (8). In mouse models of *KRAS* mutant lung cancer, MEK inhibitors display strong anti-tumor activity (15, 16). Despite these promising pre-clinical data, MEK inhibitors have failed to demonstrate efficacy in patients. In separate phase II and III trials, treatment with MEK inhibitors did not result in significant improvement in response rates or survival compared to standard chemotherapy in patients with *KRAS* mutant lung cancer (17, 18).

Mirroring the experience with other targeted therapies, resistance is a major limitation of MEK inhibition in the clinical setting. Studies have discovered several intrinsic mechanisms of resistance to MEK inhibitors, defined as resistance observed at the initiation of treatment. These include increased AKT signaling to bypass inhibition of the MAPK pathway (19–21), activation of STAT3 (22, 23), induction of ERBB3 (24) and KRAS dimerization (25), all of which may contribute to the low objective response rate observed in KRAS mutant NSCLC (17, 18). While intrinsic resistance is well examined, acquired resistance to MEK inhibitors, defined as resistance that develops in patients that initially respond to therapy, remains less understood, with ERK reactivation by FGFR upregulation the best characterized mechanism described to date (26).

Trametinib was initially discovered to induce cell cycle arrest in colorectal cancer cell lines in a *RB1* dependent manner (27). Additionally, recent findings in lung cancer have found that RB1 and p16/CDKN2A are activated by trametinib, and have implicated RB status in sensitivity to MEK inhibitors in KRAS mutant lung cancer cells; however, the underlying processes responsible for this observation remain poorly understood (28, 29). Understanding both intrinsic and acquired resistance to MEK inhibitors will be essential for defining effective clinical strategies that employ MEK inhibitors in KRAS mutant lung and other cancers, and improving overall patient outcomes.

Here, we investigated acquired resistance to MEK inhibition by generating isogenic pairs of trametinib-sensitive and – resistant KRAS mutant lung cancer cell lines through trametinib dose escalation studies. To our knowledge, these represent the first reported models of acquired resistance to MEK-targeted agents in KRAS mutant lung cancer, affording a unique opportunity to investigate genetic mechanisms of resistance in this important clinical context. Through targeted DNA sequencing, we identified mutations associated with resistance and assessed the impact of *RB* loss via CRISPR-mediated genetic knockout. Importantly, we characterize a paradoxical “drug-addicted” state in one of our models where survival is dependent on sustained MEK inhibition and demonstrate that amplification of the *KRAS*-mutant allele mediates toxicity. This work provides insight towards better understanding trametinib resistance and improving the clinical utilization of MEK inhibitors for the treatment of patients with KRAS mutant lung cancer.

## Materials and Methods

### Cells lines and reagents

All cells were cultured at 37°; air; 95%; CO2, 5%. H358 (NCI-H358), H23 (NCI-H23), H1792 (NCI-H1792) and 293T cells were obtained from American Type Tissue Culture (ATCC). Cells were regularly checked for mycoplasma contamination by polymerase chain reaction (30) and found to be negative. H358 sg*RB1*#4 parental and resistant cell lines were verified by STR profiling (Labcorp, Burlington, NC, USA). LUAD cells were grown in RPMI-1640 medium (Gibco, 11875119) supplemented with 10% fetal bovine serum (FBS) (Gibco, 12483020) and 1% Pen/Strep (Gibco, 15140-122). 293T cells were grown in DMEM medium complete with 10% FBS and 1% Pen/Strep (Gibco, 15140-122). For cells and experiments with doxycycline-inducible constructs, cells were grown in RPMI-1640 medium (Gibco, 11875119) supplemented with 10% tetracycline-free FBS (Clontech, 631101) and Pen/Strep (Gibco, 15140-122). Doxycycline hyclate (Sigma-Aldrich, D9891) was added to cells at 200 ng/mL when indicated. Trametinib (Selleckchem, S2673), SCH772984 (Selleckchem, S7101), AMG 510 (Selleckchem, S8830), SB 747651A (Tocris, 4630), MK-2206 (Selleckchem, S1078), NSC 23766 (Selleckchem, S8031), dabrafenib (Selleckchem, S8031), infigratinib (Selleckchem, S2183) and N-acetyl-L-cysteine (Sigma-Aldrich, A7250) were added to cells when indicated. Experiments were performed on cells between passages 4-20.

### CRISPR/Cas9 modification

The sgRNA sequence for *RB1* (5’-GCTCTGGGTCCTCCTCAGGA-3’) was cloned into lentiCRISPRv2 (Addgene #52961) plasmid and the co-transfected with psPAX2 (Addgene #12260) and pMD2.G (Addgene #12259) into 293T cells with Lipofectamine 2000 (Life Technologies, 11668019) to generate lentiviral particles. Empty lentiCRISPRv2 without sgRNA was used as control for *RB1* guide during lentivirus infection and later studies. H1792 and H358 cells were infected with viral supernatant and then selected with puromycin (Sigma-Aldrich, 540222) to generate stable lines. Single cell-derived clonal cells and polyclonal cells were established after *RB1* knockout. H358 sg*RB1*#3, H358 sg*RB1*#4, H1792 sg*RB1*#7 and H1792 sg*RB1*#14 displayed the best *RB1* knockout and were selected for continued studies along with an empty vector control for each cell line.

### Plasmids and generations of stable cell lines

pBABE GFP was a gift from William Hahn (Addgene #10668). GFP was subcloned into pENTR/D-TOPO (Invitrogen, K240020). pDONR223_KRAS_p.G12C was a gift from Jesse Boehm, William Hahn and David Root (Addgene #8166, (31)). GFP and KRAS^G12C^ were cloned by Gateway LR Clonase II enzyme mix (Life Technologies, 11791020) into pInducer20 (gift from Stephen Elledge, Addgene # 44012 (32)). The custom *RB1* construct was ordered from Twist Biosciences (See supplemental for full sequence). The custom sequence was printed directly into a Twist Cloning Vector, and was directly cloned into pInducer20 by Gateway LR Clonase II enzyme mix. Lentivirus was generated by transfecting 239T cells with psPAX2 (Addgene #12260) and pMD2.G (Addgene #12259) and according expression vector with Lipofectamine 2000 (Life Technologies, 11668019). H358, H23, H1792 and H358 sg*RB1*#4^tramR^ cells were infected with lentivirus and selected with 500 µg/mL G418 (Gibco, 10131027) for 2 weeks. Cells expressing GFP or KRAS^G12C^ were maintained as polyclonal populations.

### Generation of trametinib-resistant cells

To generate trametinib-resistant cell lines, we cultured H358 and H1792 single cell clones in trametinib starting at 10 nM or 30 nM for H1792 and H358 cells, respectively, and ending with 1 μM. Trametinib-containing media was refreshed every 2 or 3 days. Resistant cells were maintained as single cell-derived clones under constant exposure to the drugs. No vehicle-treated cell control was maintained in parallel.

### RNA interference

5 × 10^5^ cells were transfected with ON-TARGETplus siRNA pools (Dharmacon) targeting *MAPK3* (L-003592-00), *MAPK1* (L-003555-00), *KRAS* (L-005069-00-0005) or a non-targeting control (D-001810-10) at concentrations of 50 nM with DharmaFECT 1 transfection reagent (Dharmacon, T-2001-03). Cells were cultured for 48 hours after transfection and before subsequent analysis.

### Immunoblotting

Cells were harvested and lysed in RIPA buffer (G-Biosciences, CA95029-284) complete with protease/phosphatase inhibitor cocktail (Thermo, PI78446). Lysates were sonicated and protein concentration was determined by BCA protein assay kit (Pierce Protein Biology Products, 23225). Samples were denatured by boiling for 5 min in 4X loading buffer (Thermo Scientific, NP0008). Lysates were loaded on 4-12% Bis-Tris NuPage Protein Gels (NuPage, NP0336BOX), run in MOPS SDS buffer (NuPage, NP000102), transferred to PVDF Immobilon (Millipore, IPVH00010), and blocked in tris-buffered saline (BioRad, 170-6435) supplemented with 0.1% Tween20 (Fisher Scientific, BP337-500) (TBST) and 5% milk. Membranes were incubated in primary antibodies (1:1000) overnight at 4° in 5% bovine serum albumin (BSA) (Sigma, A9647-100G), washed with TBST, and then incubated in HRP-linked secondary anti-mouse or anti-rabbit (1:15000) (CST, 7076S and 7074S respectively) in 2.5% BSA for 1 hour at room temperature. The following antibodies were obtained from CST: pERK (9101S), ERK (4695S), pAKT (4060L), AKT (4691L), p-mTOR (5536S), mTOR (2983S), pFGFR Y653/654 (3471S), FGFR1 (9740S), pErbB3 Y1289 (4791S), ErbB3 (12708S), cleaved PARP (5625S), cleaved caspase 3 (9661S), cleaved caspase 7 (9491S), pH2AX (2577S), Rac1 (8631), cRAF (9422S), MEK1/2 (9122S), RAS (8955S), RB (9309S), E-cadherin (3195S), N-cadherin (13116S), vimentin (5741S), Snail (3879S), Slug (9585S), BiP (3183S), CHOP (2895S), ATF4 (11815S), p-eIF2A (9721S), pJNK (4668S), Elk1 (9182S), c-Fos (9F6) (2250S), c-Myc (D84C12) (5605S), RSK1/RSK2/RSK3 (D7A2H) (14813S), Phospho-p90RSK S380 (D3H11) (11989S), c-Jun (60A8) (9165S), Phospho-c-Jun S73 (D47G9) (3270S), FRA1 (D80B4) (5281S), p27 Kip1 (D69C12) (3686T), p21 Waf1/Cip1 (12D1) (2947S), p16 INK4A (D7C1M) (80772S) & vinculin (E1E9V) (13901S). GAPDH (sc-47724) was obtained from Santa Cruz Biotechnology. TTF1 (MA5-16406) was obtained from ThermoScientific. Densitometry was performed using FIJI software (33).

### Measurement of cell viability

To assess IC50s to trametinib, H358 and H1792 clones were seeded in 96-well plates at 5000 or 1500 cells per well, respectively, on day 0. On day 1, trametinib was added at the indicated concentrations. Seventy-two hours following trametinib treatment, cell viability was assessed by incubation in 10% alamarBlue viability dye (Life Technologies, Dal1100) for 2 hours. Absorbance was measured using a Cytation 3 Multi Modal Reader with Gen5 software (BioTek). For experiments involving doxycycline inducible constructs, H358, H23 and H1792 tetO GFP or KRAS^G12C^ cells were seeded at 6000 cells per well in a 6-well plate. H358 sg*RB1*#4^tramR^ tetO GFP and *RB1* cells were seeded at 5000 cells per well in a 6-well plate. Doxycycline (200 ng/mL), trametinib or AMG 510 was added at the time of seeding. Media was changed on day 3 and on day 7. On day 9, alamarBlue cell viability agent was added to the media at 10%. Absorbance was measured using a Cytation 3 Multi Modal Reader with Gen5 software (BioTek).

For proliferation assay, cells were seeded at 1 × 10^5^ cells per well. Trametinib was added at the time of seeding. Media was changed on day 3. On day 7, media was aspirated, and cells were washed with PBS. A 0.5% crystal violet (Sigma, HT90132), 20% methanol solution was added to cells. Cells were incubated with rocking for 15 minutes, after which crystal violet was discarded and plates were left to dry overnight.

#### Clonogenic assays

H358 sg*RB1*#4^tramR^ cells were seeded at 100 cells per well in a 6-well plate in either 0.1 % DMSO or 1 µM trametinib and propagated for 11 days. Trametinib or DMSO was refreshed every 3 days. At endpoint, media was washed out and cells were stained with crystal violet. Colonies in the scanned images of the crystal violet stained plates were quantified using FIJI software (33). Briefly, colonies on the plate were identified using the “Color Threshold” and “Watershed” commands. Identified particles were subsequently counted using the “Analyze Particles…” function (Size filter = 5-infinity, circularity filter = 0.5-1.0).

### IncuCyte growth assays

Cells were seeded at 5000 cells per well in a clear bottom 96-well plate and treated with drugs at the indicated concentrations on day 0. On day 1, cells were placed in an IncuCyte S3 live-cell imaging system contained in an incubator kept at 37°C and 5% CO_2_. Images were taken at a 4-hour intervals in quadruplicate for 120 hours. For experiments with nuclei quantification, cells were treated with Incucyte® Nuclight Rapid Red Dye for Live-Cell Nuclear Labeling (Sartorius, 4717) at time of experiment seeding for a final concentration of 1:750. For experiments with siRNA, cells were cultured for 48 hours after siRNA transfection before being seeded into a 96-well plate and placed in the IncuCyte imaging system. Cells were imaged for 136 hours.

### MSK-IMPACT sequencing

We extracted DNA from trametinib-resistant clones and their parental counterparts using a DNeasy Blood & Tissue Kit (Qiagen, 69506). DNA was submitted for profiling on the MSK-IMPACT (Integrated Mutation Profiling of Actionable Cancer Targets) platform, a hybridization capture-based next generation sequencing (NGS) platform for targeted deep sequencing of exons and selected introns from 468 cancer-associated genes and selected gene fusions (34). The assay detects mutations and copy-number alterations in samples. We compared resistant cells to their parental controls, and considered alterations detected only in the resistant cells as potential genes associated with resistance to trametinib.

#### Quantitative RT-PCR

Cells were lysed and RNA was extracted using RNeasy Mini kit (Qiagen, 74106) according to manufacturer’s protocol. cDNA was prepared using the High-Capacity RNA-to-cDNA™ Kit (Applied Biosystems, 4387406). RT-PCR reactions were performed using the TaqMan Gene Expression Master Mix (Thermo Fisher, 4369016). The following TaqMan Gene Expression Assays primers were obtained from Thermo Scientific: KRAS (Hs00364284_g1, 4331182), NRAS (Hs00180035_m1 S, 4331182), HRAS (Hs00978051_g1, 4331182) and β Actin (4333762F). Reactions were performed on an Applied Biosystems 7500 Fast Real-Time PCR System (Thermo Fisher). Relative expression was quantified using the ΔΔCt method and using the average cycle threshold.

### RAS-GTP pulldown

Cells were treated with either 0.1% DMSO or 1 µM trametinib in media containing 10% FBS for 24 hours. Cells were harvested, lysed and active RAS levels were measured by affinity purification using an Active Ras Detection Kit (Cell Signaling Technologies, 8821S). Pulldown samples were loaded on a 4-12% Bis-Tris NuPage Protein gel (NuPage, NP0336BOX), and immunoblotted using the anti-RAS antibody provided with the kit.

### Statistical analysis

Statistical analyses were performed using GraphPad Prism version 8.2.1 (GraphPad Software, San Diego, CA, USA). Nonlinear regression with fitting by least squares method was performed to determine IC50 (nM) and growth rate constant k (hours^-1^). Mean and profile likelihood 95% CI are reported. Parameters calculated for treatment conditions were compared to control parameters by Extra sum-of-squares F test. Differences in continuous variables were evaluated with a two-sided student’s *t* test. *P* values < 0.05 were considered statistically significant, indicated as following; *p < 0.05, **p < 0.01, ***p < 0.001, ****p < 0.0001, NS = not significant.

## Results

### Establishment of KRAS-Mutant Lung Adenocarcinoma Cells Demonstrating Acquired Resistance to Trametinib

Acquired resistance to MEK inhibition in lung cancer has previously been associated with p16/RB1/CDK4 regulatory status (28, 29). Thus, in order to model this scenario in KRAS mutant LUAD, we first generated isogenic clones of H358 and H1792 cell lines with CRISPR/Cas9-mediated *RB1* knockout. H358 and H1792 both bear KRAS^G12C^ activating mutations and are highly dependent on signaling through the RAS-RAF-MEK-ERK pathway for survival. Two single cell derived clones from H358 (H358 sg*RB1*#3 and H358 sg*RB1*#4) and H1792 (H1792 sg*RB1*#7 and H1792 sg*RB1#*14) were chosen based on the degree of *RB1* knockout displayed. An empty vector control cell line was also established for each cell line (Supplemental Figure 1A). Dose response curves and IC50 values were calculated for all cell lines by non-linear regression with fitting by least squares method, and demonstrate that all clones were sensitive to low doses of trametinib (H1792 sgControl = 20.3 nM, 95% CI 14.4-28.5; H1792 sg*RB1*#7 = 22.4 nM, 95% CI 16.4-30.4; H1792 sg*RB1*#14 = 37.6 nM, 95% CI 28.6-49.4; H358 sgControl = 3.7 nM, 95% CI 2.7-5.1; H358 sg*RB1*#3 = 5.9 nM, 95% CI 3.9-8.9; and H358 sg*RB1*#4 = 6.8 nM, 95% CI 5.1-9.1; Figure 1A and 1B, Table 1). IC50s calculated for *RB1* KO clones were compared to the IC50 calculated for the control clone by extra sum-of-squares F test. In H1792 clones, *RB1* knockout resulted in a modest increase in IC50 for *RB1* KO clones, which was found to be significant in H1792 sg*RB1*#14 (p<0.0001) when compared by extra-sum-of-squares F test. In both H358 clones, *RB1* knockout resulted in modest but consistent increases in trametinib IC50 relative to the vector control, which was significant for both clones (p=0.0305 and p=0.0002 for H358 sg*RB1*#3 and H358 sg*RB1*#4 respectfully). This is consistent with previous observations of *RB1* loss decreasing sensitivity to trametinib in H358 cells (28, 29). Each clonally expanded cell line was treated with escalating doses of trametinib until they were able to consistently grow in a concentration of 1 µM, at which point they were considered resistant. We observed no difference in the rate at which *RB1* KO or control clones acquired resistance (Supplemental Figure 1B). The IC50s for inhibition of growth by trametinib were then re-assessed and found to be > 10 µM for each resistant clone (referred to from this point with “^tramR^” after the cell line name) (Figure 1C and 1D, Table 1). Growth of parental cells was inhibited when cultured in 1 µM trametinib over a 5-day period (Supplemental Figure 1C-H), while in contrast, resistant clones proliferated under these conditions (Supplemental Figure 1C-H). H358 sg*RB1*#3^tramR^ and H358 sg*RB1*#4^tramR^ display faster growth in 1 µM trametinib than their parental counterparts in 0.1% DMSO (Supplemental Figure 1D and 1E). All other resistant clones grow at relatively similar rates to their parental counterparts in the absence of drug. Of note, dose-response assays on H358 sg*RB1*#4^tramR^ produce a bell-shaped curve, suggesting these cells are more viable when grown in a certain range of drug concentration than when grown in 0.1% DMSO (Figure 1C).

**Figure 1.**
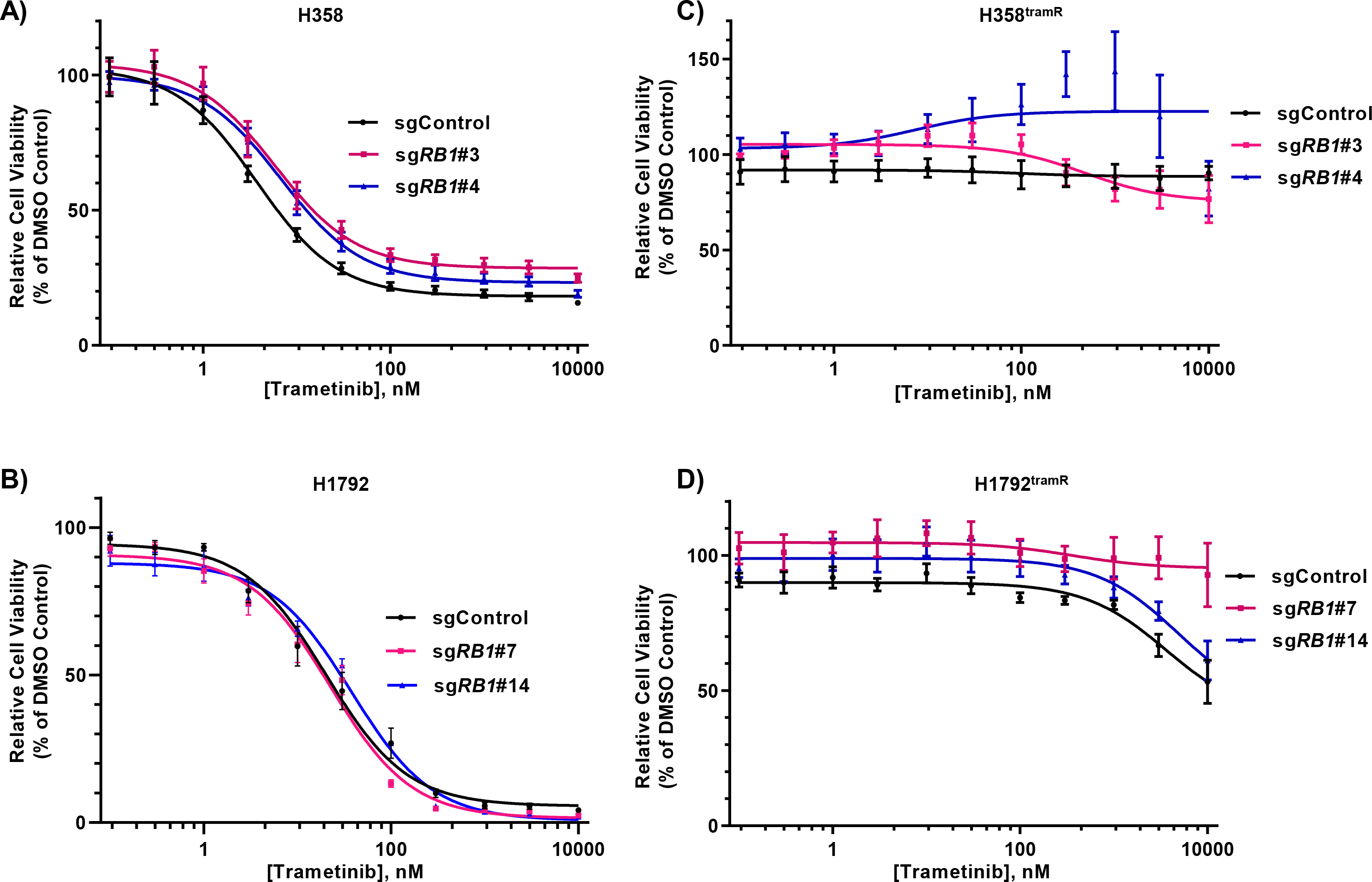
Impact of RB1 on trametinib resistance. **a, b. Isogenic H358 and H1972 clones with** RB knockout were grown in the indicated concentrations of trametinib (1 nM to 10 µM) for 3 days. Cell viability was assayed with alamarBlue and relative viability was calculated as a percent of the 0.1% DMSO-treated control. Error bars are SEM from 3 independent experiments. **c, d.** Resistant RB knockout and control H358 and H1792 clones were grown in the indicated concentrations of trametinib (1 nM to 10 µM) for 3 days. Cell viability was assayed with alamarBlue and viability was calculated relative to 0.1% DMSO vehicle control. Error bars are SEM from 3 independent experiments.

**Table 1.** Summary table of calculated IC50 values for parental and trametinib resistant cell lines.

| **Cell Line** | **Parental IC<sub>50</sub> nM** | **TramR IC<sub>50</sub> μM** |
| --- | --- | --- |
| H1792 sgControl | 20.3 (14.4-28.5) | > 10 |
| H1792 sgRB1#7 | 22.4 (16.4-30.4) | > 10 |
| H1792 sgRB1#14 | 37.6 (28.6-49.4) | > 10 |
| H358 sgControl | 3.7 (2.7-5.1) | > 10 |
| H358 sgRB1#3 | 5.9 (3.9-8.9) | > 10 |
| H358 sgRB1#4 | 6.8 (5.1-9.1) | > 10 |

To assess the status of KRAS-related signaling pathways and previously reported mechanisms of MEK inhibitor resistance in the trametinib resistant cells, we performed immunoblot analysis of key downstream effectors. All resistant clones – except for H358 sg*RB1*#4^tramR^ – displayed dramatically decreased phospho-ERK in comparison to their parental counterparts, suggesting that these cell lines have bypassed the requirement for the MAPK signaling pathway for growth (Supplemental Figure 1J). H358 sg*RB1*#3^tramR^, H358 sg*RB1*#4^tramR^ and H1792 sg*RB1*#14^tramR^ display increased pAKT levels, indicating that PI3K/AKT activation, a common mechanisms of adaptive resistance to MEK inhibition, may compensate for diminished MAPK activity and mediate cell survival in the presence of trametinib (19). ERBB3 is also upregulated in all three H358^tramR^ clones (Supplemental Figure 1J) and has previously been shown to activate PI3K signaling and drive resistance to targeted therapy (24). Lastly, increased expression of FGFR1 due to feedback inhibition has been reported to induce resistance to MEK inhibition (26), and was observed in H358 sgControl^tramR^ and H358 sg*RB1*#4^tramR^ cells (Supplemental Figure 1J). Activation of epithelial to mesenchymal transition (EMT) genes has also been reported in cases of resistance to targeted therapies in lung cancer (35–37). Treatment naïve H358 and H1792 cells have differing expression of EMT genes, with the latter being more mesenchymal like, which may influence mechanisms of resistance. H358 sg*RB1*#4^tramR^ also displayed upregulation of N-cadherin, vimentin, snail and slug as well as downregulation of E-cadherin (Supplemental Figure 1J), all of which are associated with an FGFR1-regulated mesenchymal-like state in KRAS mutant LUAD (37). This suggests that H358 sg*RB1*#4^tramR^ may have undergone EMT while developing resistance to trametinib. Images of parental and resistant H358 sg*RB1*#4 cells also suggest a morphological shift to a more mesenchymal-like phenotype (Supplemental Figure 1I). However, the lack of phosphorylated FGFR1 indicates that the cells may not be reliant on FGFR1 signaling for survival. Overall, these data suggest that the trametinib-resistant cell lines have bypassed the requirement for MAPK pathway signaling, and instead may rely on activated ERBB3-PI3K-AKT pathways to sustain cancer cell survival in the face of MEK inhibition.

### Drug Removal Leads to Cell Death in Selected Trametinib-Resistant Lung Cancer Cells

Assessment of known mechanisms of resistance to MEK inhibitors offered potential insights into the processes driving acquired resistance in our isogenic model systems. Upon further characterization, we found that H358 sg*RB1*#4^tramR^ was *dependent* on continued culture in trametinib for survival. Using the IncuCyte S3 live-cell imaging system, we measured well confluence and nuclei count over time. We performed logistic growth regression using confluence measurements to determine the growth rate of cells under different treatments with calculated growth rates compared by extra sum-of-squares F test. Paradoxically, H358 sg*RB1*#4^tramR^ cells have a significantly higher growth rate (p<0.0001) when cultured in 1 µM trametinib (0.03734 h^-1^, 95% CI 0.03547 h^-1^-0.03927 h^-1^) than when the drug is withdrawn (0.02395 h^-1^, 95% CI 0.02106 h^-1^-0.02692 h^-1^), in contrast to the parental counterpart (Figure 2A, Supplemental Figure 2A). This relationship is also seen when assessing nuclei counts (Supplemental Figure 2C). H358 sg*RB1*#4^tramR^ cells also display significantly poorer colony forming ability relative to their parental counterpart (Figure 2B). While parental H358 sg*RB1*#4 cells can proliferate in 0.1% DMSO but are inhibited by 1 µM trametinib, H358 sg*RB1*#4^tramR^ only grow in 1 µM trametinib and not in 0.1% DMSO (Figure 2C). Together, this suggests that the cells – which were initially sensitive to trametinib – have subsequently become “addicted” to the drug in the process of acquiring resistance. When H358 sg*RB1*#4^tramR^ is grown without trametinib, cells develop vacuoles, similar to the phenotype we have previously reported that coincides with hyperactive MAPK signaling in KRAS mutant lung cancer cells (38) (Figure 2D). Bright field images confirmed the increased proliferation in 1 µM trametinib, as well as appearance of vacuoles around 72 hours following drug removal (Supplemental Figure 2A). Withdrawal of trametinib also corresponds to activation of caspases 3 and 7, as well as PARP cleavage, suggesting that drug removal induces apoptosis (Figure 2E). The drug addiction phenotype was only observed in H358 sg*RB1*#4^tramR^, with all other resistant clones demonstrating no adverse effects when trametinib was removed. Parental and resistant H358 sg*RB1*#4 cells were submitted for STR profiling and were confirmed to be H358 cells (Supplemental Figure 2B). Interestingly, H358 sg*RB1*#4^tramR^ was the only trametinib-resistant clone with appreciable levels of pERK (Supplemental Figure 1J), suggesting activation of this pathway may play a role in mediating the drug-addicted state.

**Figure 2.**
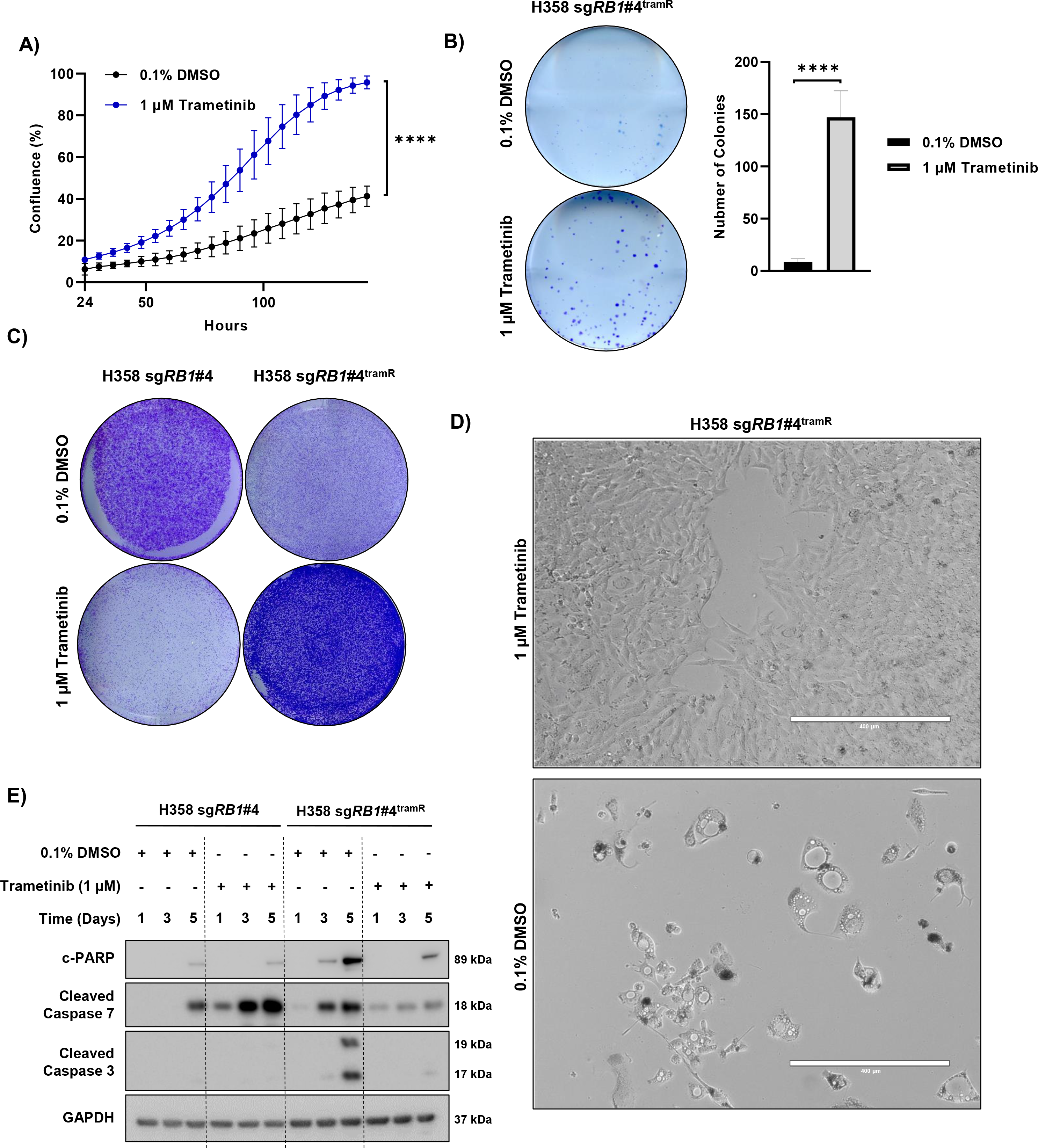
*H358* sgRB1#4^tramR^ cells are addicted to trametinib. **a.** H358 sg*RB1*#4^tramR^ grow slower in 0.1% DMSO then in 1 µM trametinib as measured by IncuCyte S3 live-cell imaging system. Error bars are 95% confidence interval from 4 independent experiments. P value from extra sum-of-squares F test on calculated logistic growth rate are indicated. ****p < 0.0001. **b** Clonogenic growth assay performed on H358 sg*RB1*#4^tramR^ grown in 0.1% DMSO or 1 µM trametinib. Cells were fixed and stained with crystal violet following 14-day treatment under the indicated conditions. Representative images from 4 independent replicates. Colonies were quantified using Fiji. P value from students t test on colony number shown, ****p < 0.0001. **c.** H358 sg*RB1*#4^tramR^ cells were grown in either 0.1% DMSO or 1 µM trametinib for 7 days, then stained with crystal violet. H358 sg*RB1*#4^tramR^ can proliferate better in 1 µM trametinib than in 0.1% DMSO vehicle, the opposite of what is seen in their parental counterparts. **d** 10X microscope images were taken after 11 days. Vacuoles form in H358 sg*RB1*#4^tramR^ cells when grown without trametinib. Scale bar shown represents 400 µm. **e** H358 sg*RB1*#4 parental and resistant cells were grown in either 0.1% DMSO or 1 µM trametinib and harvested after 1, 3 or 5 days. Lysates were subjected to immunoblotting for the indicated proteins. H358 sg*RB1*#4^tramR^ cells display upregulation of apoptosis markers when grown without trametinib.

### Addiction to MEK Inhibitor Treatment is Mediated by ERK2

Our observation that MEK inhibitor withdrawal leads to cancer cell death mirrors similar reports of targeted therapy addiction in melanoma (39–43), lung cancer (44) and lymphoma (45). In these reports, resistant cells have become dependent on suppression of the MAPK signaling pathway for survival, implicating hyperactivation of the MAPK pathway – and in some instances hyper-phosphorylation of ERK2 specifically – as the driver of the drug addiction phenotype. Our previous work has demonstrated that hyperactivation of ERK2 is toxic to lung adenocarcinoma cells bearing KRAS or EGFR oncogenic mutations (46). Given that H358 sg*RB1*#4^tramR^ demonstrates addiction to a MEK inhibitor, and that MEK1/2 directly activates ERK1/2, we next assessed the effects of drug withdrawal on ERK1/2 phosphorylation. We observed that removal of trametinib corresponds to a major rebound in pERK levels within 30 minutes and persists past 72 hours, decreasing with time (Figure 3A). Trametinib removal also corresponds with an increase in downstream targets of pERK including pRSK after 30 min, increases in cFOS and p-cJun after 1 hour and upregulation of FRA1 after 3 hours (Supplemental Figure 3A). While p-cJun and cFOS increases appear to be transient, pRSK and FRA1 increases are sustained past 72 hours. Markers of apoptosis are also induced after the pERK increase, 48 hours after drug removal, and coinciding with pH2AX, a marker of double stranded DNA breaks (Figure 3A). We investigated makers of ER stress in H358 sg*RB1*#4^tramR^ and found that removal of trametinib results in upregulation of BiP, a chaperone protein upregulated in response to unfolded proteins in the ER, and of CHOP, a transcription factor known to activate apoptosis in response to ER stress (Supplemental Figure 3B). CHOP is upregulated 12 hours after drug removal along with ATF4 and p-eIF-2A, two activators of the protein, suggesting an ER stress response may be driving subsequent apoptosis.

**Figure 3.**
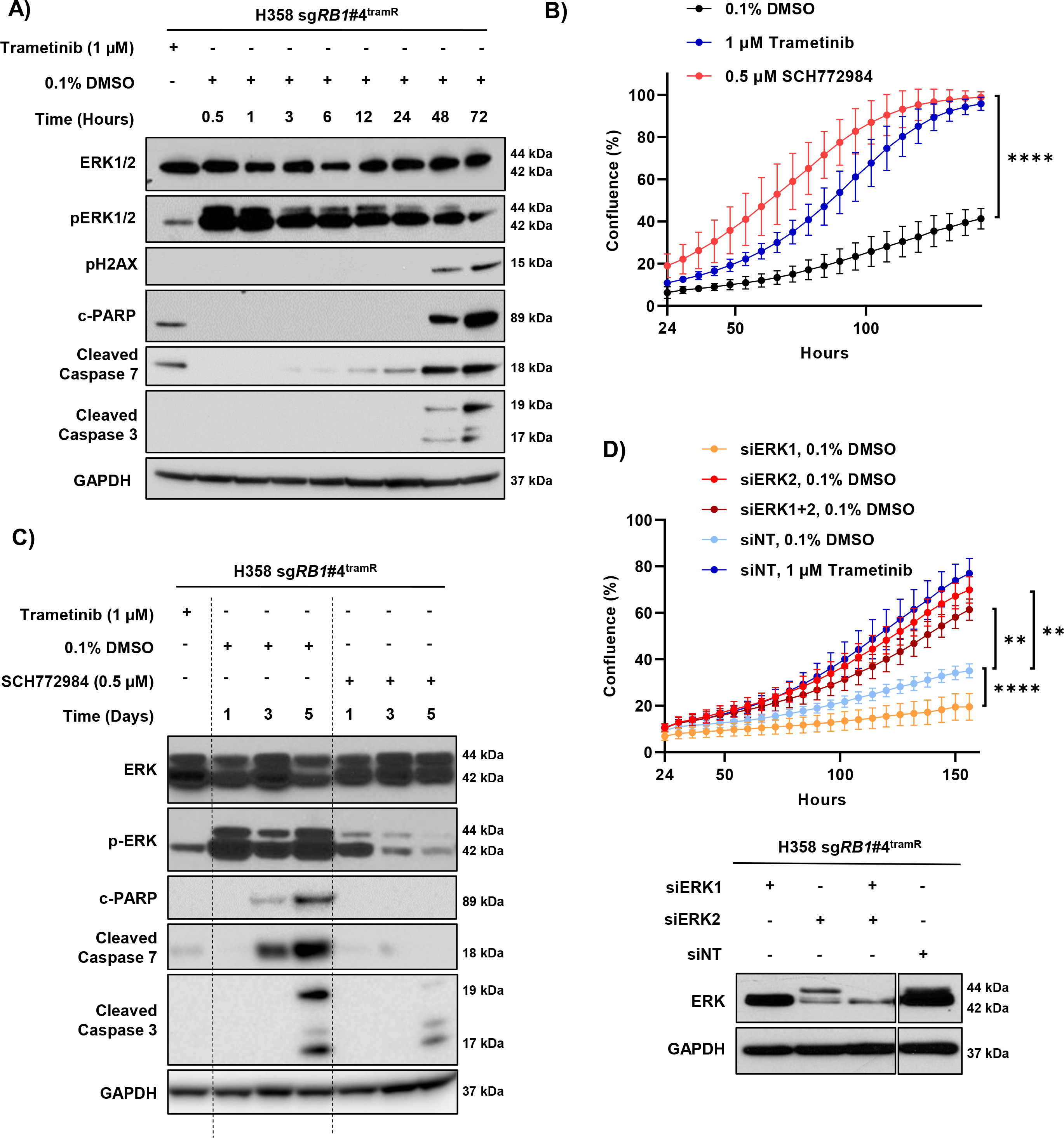
ERK2 hyperactivation mediates trametinib addiction. **a** H358 sg*RB1*#4^tramR^ were treated with 0.1% DMSO or 1µM trametinib, harvested after the indicated time periods, and immunoblotted for the proteins shown. Starting at 30 min after drug removal, and persisting past 72 hours, there is a strong pERK rebound, as well as induction of markers of apoptosis and DNA damage. **b** H358 sg*RB1*#4^tramR^ cells were seeded in the indicated concentrations. Inhibition of ERK with 0.5 µM SCH772984 rescues H358 sg*RB1*#4^tramR^ cell growth after trametinib removal, as measured by IncuCyte S3 live-cell imaging system. Error bars are 95% confidence interval from 4 independent replicates. P value from extra sum-of-squares F test on calculated logistic growth rate is indicated. ****P< 0.0001. **c** H358 sg*RB1*#4^tramR^ cells were treated with indicated drug concentrations for indicated time, harvested, lysed and immunoblotted. Treatment with 0.5 µM SCH772984 rescues induction of pERK and apoptosis markers. **d.** siRNA targeting ERK1 and/or ERK2 were transfected into H358 sg*RB1*#4^tramR^. Knockdown of ERK2 alone, or ERK1 and ERK2, rescues cells from death after trametinib removal. Knockdown of ERK1 alone further inhibits cell growth following trametinib removal. Confluence was measured by IncuCyte S3 live-cell imaging system. Error bars represent SEM from 4 independent experiments. P values from student’s t test on confluence at endpoint growth rate are indicated. **p < 0.01, ****p < 0.0001.

To validate pERK as the effector of this paradoxical drug addiction phenotype, we attempted to rescue H358 sg*RB1*#4R cells from trametinib withdrawal by treatment with SCH772984, an ERK1/2 inhibitor (47). Treatment with 0.5 µM SCH772984 results in full rescue of cell death following trametinib removal as indicated by logistic growth regression analysis (0.5 µM SCH772984 = 0.03975 h^-1^, 95% CI 0.03716 h^-1^ to 0.04244 h^-1^; 0.1% DMSO = 0.02395 h^-1^, 95% CI 0.02106 h^-1^ to 0.02692 h^-1^; p<0.0001) (Figure 3B). Treatment with SCH772984 reduces pERK to levels similar to treatment with 1 µM trametinib, highlighting the suppression of ERK hyperactivation after MEK inhibitor withdrawal (Figure 3C). SCH772984 treatment also rescues cells from induction of apoptosis markers. To assess the role of ERK2 specifically, we performed siRNA knockdown of ERK1 and ERK2 alone or in combination in the drug-addicted cells. We observed that knockdown of ERK2 rescued cell growth following trametinib removal, whereas knockdown of ERK1 alone further inhibited cell growth under this condition (Figure 3D). At endpoint, confluence of cells grown in 0.1% DMSO and treated with siERK2, siERK1+2 or siERK1 were significantly different than confluence of cells grown in 0.1% DMSO treated with siNT (p=0.0016, p=0.0023 and p<0.0001, respectively). Together, these findings demonstrate that MEK inhibitor withdrawal leads to acute hyperactivation of ERK2, which causes ER stress and subsequent apoptosis in MEK inhibitor-addicted resistant cells.

To investigate the role *RB1* may play in the drug addiction phenotype, we re-expressed *RB1* cDNA possessing silent mutations at the sgRNA binding sequence to avoid cleavage using a doxycycline inducible vector, with inducible GFP serving as a control (Supplemental Figure 3C). Induction of *RB1* expression has no effect on pERK when the cells are grown in trametinib (Supplemental Figure 3C). To assess if *RB1* affects cell proliferation, we treated cells with doxycycline and cultured them with or without 1 µM trametinib for 9 days and noted no significant change upon induction of RB1, either in the presence or absence of trametinib (Supplemental Figure 3D). Crystal violet staining reveals no differences in proliferation when *RB1* is induced either with or without 1 µM trametinib, relative to GFP control states (Supplemental Figure 3E). Together, these results suggest that *RB1* does not play a role in the drug addiction phenotype and that the drug addicted phenotype could have developed in RB proficient cells.

### Acquired Genetic Alterations in the MAPK Signaling Pathway in Drug Addicted Cells

In order to elucidate mechanisms of acquired resistance and addiction to trametinib, we performed targeted sequencing using the MSK-IMPACT panel (34). Sequencing detected CRISPR induced *RB1* mutations in the two resistant H358 clones as P28Qfs*35 and E30* for H358 sg*RB1*#3^tramR^ and H358 sg*RB1*#4^tramR^ respectively. In addition to a candidate F53V mutation identified in *MAP2K1* (encoding MEK1) that could potentially mediate resistance in H1792 sgControl^tramR^ cells (Supplemental Figure 4B), this analysis revealed copy number alterations of key MAPK regulators in H358 sg*RB1*#4^tramR^ that could potentially regulate the MEKi withdrawal phenotype. This included copy number amplification of *KRAS* and *RAF1* (encoding C-RAF), as well as gains of *MAP2K2* (encoding MEK2) and *RAC1* (Figure 4A). All of these genes have been reported to play a role in ERK activation, which we demonstrated has a crucial function in trametinib addiction. We found that H358 sg*RB1*#4^tramR^ has increased RAS, CRAF, RAC1 and MEK2 protein levels, confirming the downstream consequence of genomic amplification (Figure 4B). H358 cells are heterozygous for mutant *KRAS* with one wild type and one mutant allele. MSK-IMPACT reveals that the KRAS^G12C^ mutant allele is the one amplified in both parental and resistant H358 sg*RB1*#4. Overall, we found that the MAPK pathway is potentially activated in H358 sg*RB1*#4^tramR^ cells at three different nodes above ERK (Figure 4C), suggesting that one or more of these alterations may drive ERK hyperactivation after trametinib withdrawal.

**Figure 4.**
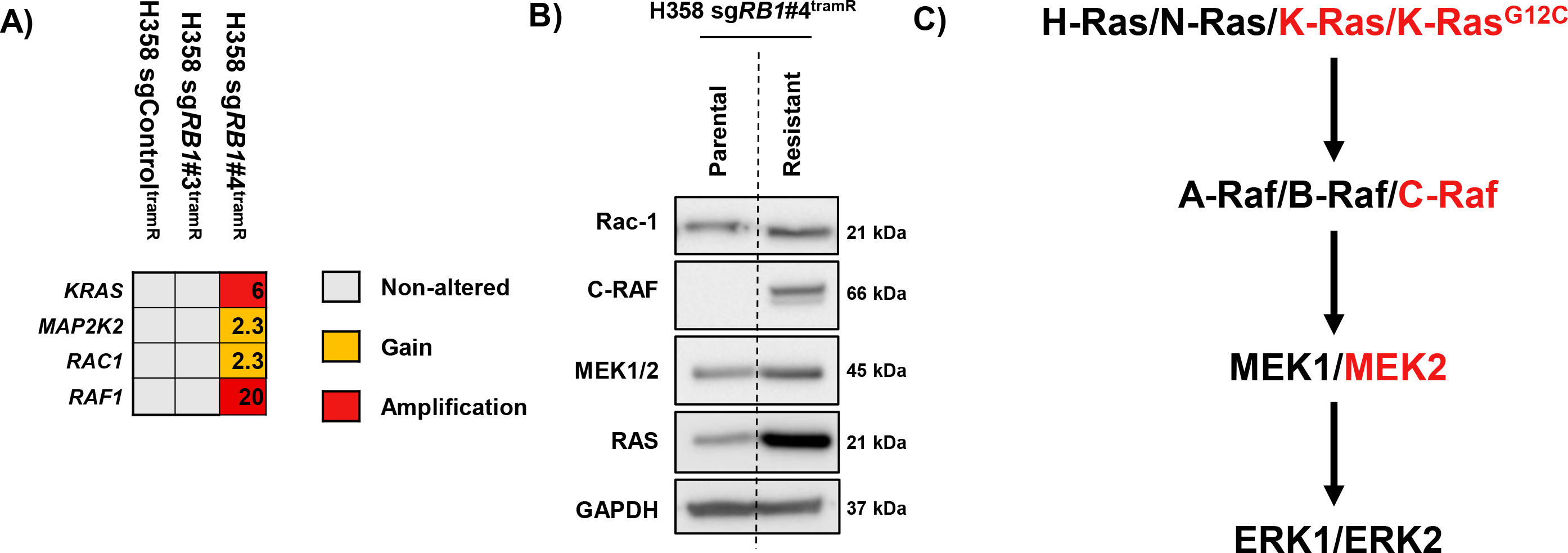
MAPK pathway components are amplified in H358 sg*RB1#4^tramR^.* **a.** MSK-IMPACT profiling reveals *RAC1*, *RAF1*, *MAP2K2* and *KRAS* amplification. Copy number alterations of *RAC1*, *RAF1*, *MAP2K2* and *KRAS* are indicated. **b.** H358 sg*RB1*#4 parental and resistant cells were cultured in 0.1% DMSO or 1 µM trametinib respectively, harvested, and immunoblotted for the indicated proteins. Genes that were amplified in (**a)** were validated at the protein level. **c.** Proteins are amplified at 3 different nodes above ERK1/2 in the MAPK pathway.

### KRAS^G12C^ Amplification Results in ERK Hyperactivation Following Trametinib Withdrawal

We have previously shown that overexpression of KRAS^G12V^ in H358 cells leads to ERK hyperactivation and cellular toxicity (46). To evaluate mutant *KRAS* amplification as a potential mediator of the drug addicted phenotype, we first compared RAS activity levels in H358 sg*RB1*#4 parental and resistant cell lines by affinity purification for active GTP-bound RAS. This revealed a major increase in RAS activity in the resistant cells (Figure 5A). We performed qPCR on both parental and tramR H358 sg*RB1*#4 cells and confirmed that KRAS is the only RAS isoform overexpressed in the resistant context (Figure 5C). Based on this observation, we hypothesized that inhibiting KRAS may circumvent the toxic effects of MEK inhibitor withdrawal in H358 sg*RB1*#4^tramR^ cells. To test this hypothesis, we knocked down KRAS with siRNAs and observed no difference in cell viability after removal of trametinib (Figure 5B). However, KRAS knockdown did re-sensitize H358 sg*RB1*#4^tramR^ to trametinib, suggesting KRAS amplification mediates trametinib resistance (Figure 5B). We rationalized that while H358 sg*RB1*#4^tramR^ are no longer as dependent on MAPK signaling as their parental counterparts, they are likely still dependent on KRAS signaling that is tuned within an appropriate level. Additionally, H358 sg*RB1*#4^tramR^ may be dependent on KRAS signaling though the AKT pathway by activation of PI3K. Thus, complete knockdown of KRAS may lead to cell death, regardless of MEK inhibition. We next sought to specifically suppress KRAS^G12C^ signaling using AMG 510, a novel small molecule inhibitor specific to the G12C form of the oncoprotein (48). By inhibiting KRAS^G12C^ with 0.5 µM AMG 510, we achieved full rescue of H358 sg*RB1*#4^tramR^ proliferation following trametinib removal, with a growth rate (0.03308 h^-1^, 95% CI 0.03131 h^-1^ to 0.03488 h^-1^) significantly higher (p<0.0001) than observed in 0.1% DMSO (0.02395 h^-1^, 95% CI 0.02106 h^-1^ to 0.02692 h^-1^) and comparable to that seen with 1 µM trametinib (Figure 5D). Similar to treatment with the ERK1/2 inhibitor SCH772984, treatment with AMG 510 also suppressed the pERK rebound following removal of trametinib and partially prevented induction of cleaved PARP, cleaved caspase 3 and cleaved caspase 7 (Figure 5E).

**Figure 5.**
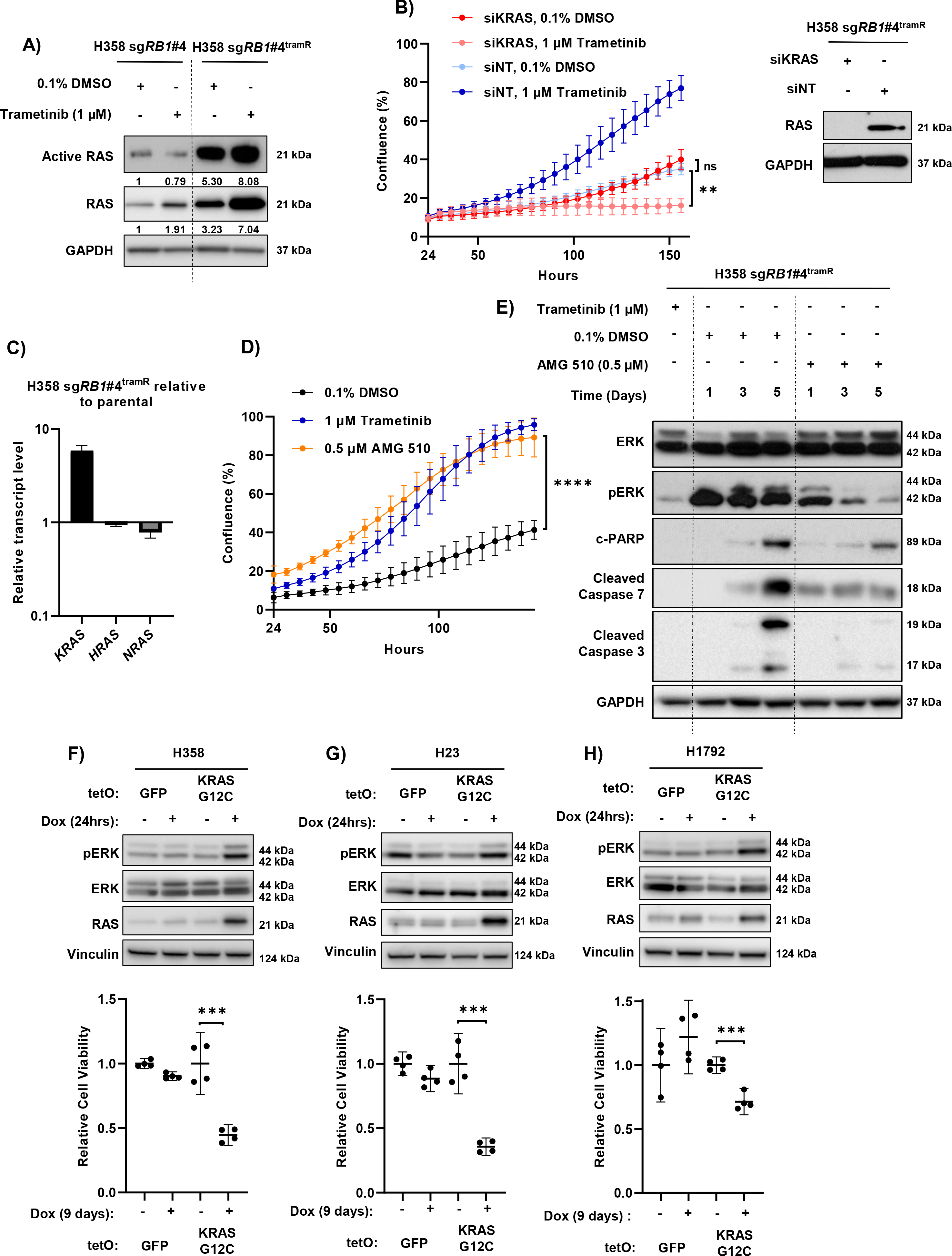
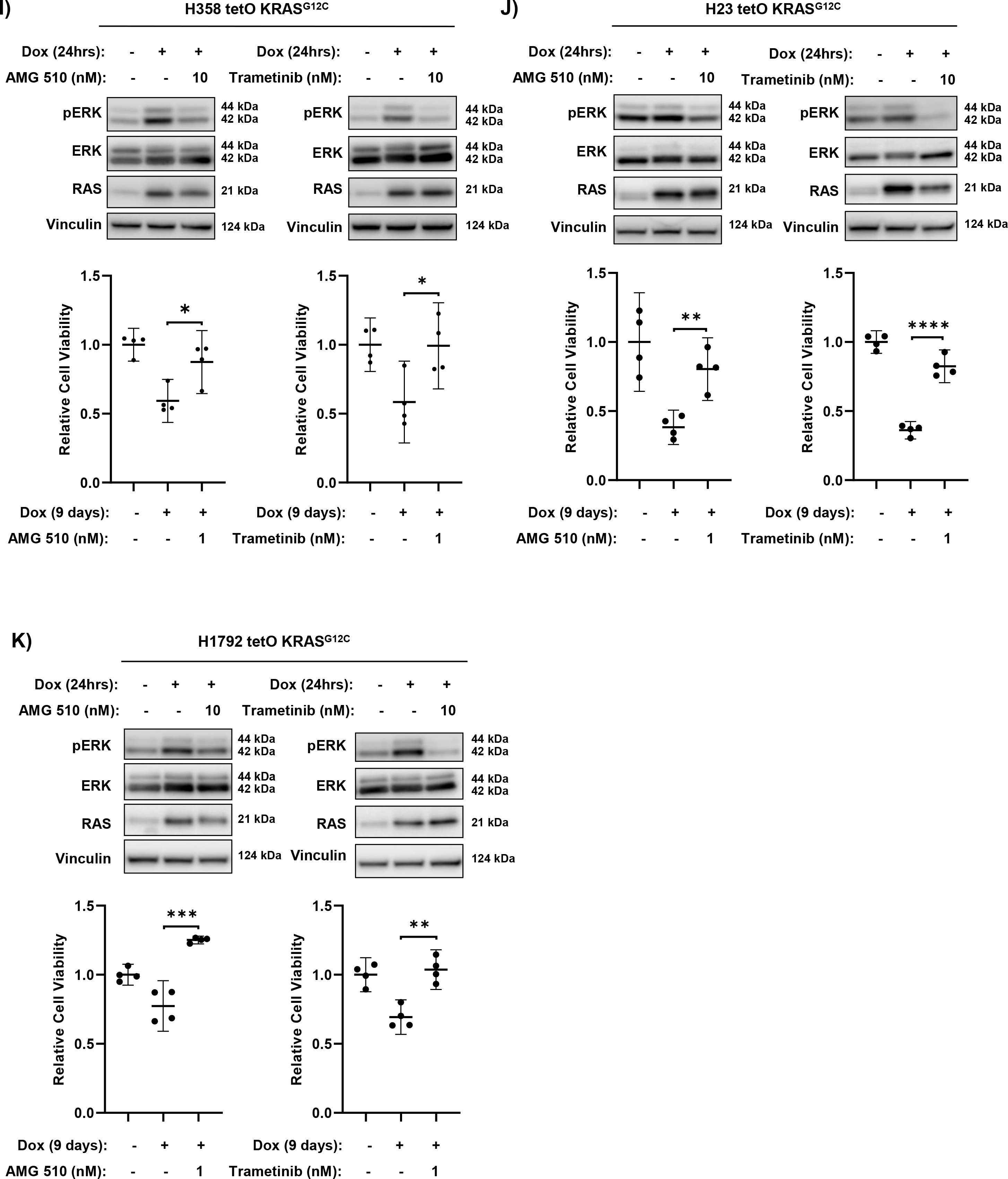
Mutant KRAS amplification drives hyperactivation of ERK and drug addiction following trametinib removal. **a.** Active GT-bound RAS was isolated by affinity purification,. H358 sg*RB1*#4^tramR^ cells have much higher levels of active RAS compared to their parental counterparts. Protein levels were quantified by densitometry using FIJI. Normalized active and total RAS levels relative to H358 sg*RB1*#4 parental treated with 0.1% DMSO are shown. **b** KRAS knockdown by siRNAs does not rescue drug addiction in H358 sg*RB1*#4^tramR^, as measured by IncuCyte S3 live-cell imaging system. The loading control used for this figure (GAPDH) is the same as the one used in Figure 3D. Error bars are SEM from 4 independent experiments. p value from student’s t test on confluence at endpoint growth rate are indicated. NS = not significant. **c** KRAS RNA levels are increased in H358 sg*RB1*#4^tramR^ cells compared to parental counterparts. **d** Inhibition of KRAS^G12C^ with 0.5 µM AMG 510 rescues H358 sg*RB1*#4^tramR^ cell growth after removal trametinib, as measured by IncuCyte S3 live-cell imaging system. Error bars are 95% confidence interval from 4 independent experiments. P value from extra sum-of-squares F test on calculated logistic growth rate is indicated. ****P< 0.0001. **e** Treatment with 0.5 µM AMG 510 partially rescues induction of pERK and apoptosis markers in H358 sg*RB1*#4^tramR^. **f, g, h** H358, H23 and H1792 were engineered to stably express KRAS^G12C^ under the control of a doxycycline inducible as described in the methods. GFP or KRAS^G12C^ expression was induced by adding 200 ng/mL doxycycline to the media for the indicated amounts of time. Induction of KRAS^G12C^ after 24 hours leads to increases in pERK levels. Cell viability measured by adding alamarBlue after 9-day treatment with doxycycline, calculated relative to no doxycycline control. Induction of KRAS^G12C^ over 9 days reduces cell viability in the 3 cell lines compared to the no doxycycline control. Error bars are 95% confidence interval from 4 independent experiments. **i, j, k** Inhibition of MEK or KRAS^G12C^ specifically with 10 nM trametinib or 10 nM AMG 510 partially rescues pERK by KRAS^G12C^ after 24 hours. After 9 days, treatment with 1 nM trametinib or 1 nM AMG 510 also partially rescues loss of cell viability driven by induction of KRAS^G12C^, as measured by alamarBlue. The error bars represent 95% confidence interval from 4 independent experiments. P values from student’s t test are indicated. *P < 0.05, **p < 0.01, ***p < 0.001, ****p < 0.0001, NS = not significant.

To rule out the involvement of other pathways in regulating the trametinib addiction phenotype, we performed similar experiments attempting to rescue H358 sg*RB1*#4^tramR^ cells from trametinib withdrawal by inhibiting other proteins that are amplified upon the acquisition of resistance, or pathways previously implicated in reports of drug addiction. In the only previous report of drug addiction in lung cancer cells, the authors demonstrate the rescue of this phenotype with AKT inhibition (44). To test this in our model, we attempted to rescue H358 sg*RB1*#4^tramR^ cells with an AKT inhibitor, MK-2206 (49), but found no effect (Supplemental Figure 5A). FGFR1 was also found to be upregulated in in H358 sg*RB1*#4^tramR^, however treatment with the FGFR1 inhibitor infigratinib (50) did not rescue cell death following trametinib removal (Supplemental Figure 5B). Indeed, higher concentrations of infigratinib inhibited proliferation following drug removal and points to FGFR1 upregulation mediating trametinib resistance in this cell line, but not trametinib dependence. We also attempted to rescue the drug-addicted phenotype with dabrafenib (51) and NSC 23766 (52), inhibitors of CRAF and RAC1, respectively, which, like KRAS, were amplified in H358 sg*RB1*#4^tramR^ cells. However, as with AKT and FGFR1 inhibition, these inhibitors could not circumvent cell death after trametinib removal at any concentration tested (Supplemental Figure 5C and 5D).

To validate KRAS^G12C^ amplification as the determinant of ERK hyperactivation and cellular toxicity in the absence of MEK inhibition, we introduced exogenous KRAS^G12C^ under the control of a doxycycline inducible promoter into H358, H23 and H1792 cells, which all harbor a single endogenous mutant allele of KRAS^G12C^. Stable polyclonal populations of H358, H23 and H1792 were created by lentiviral infection and subsequent selection. Mutant KRAS or GFP control, were subsequently induced by adding doxycycline to culture media. In H358, H23 and H1792 cells, induction of exogenous KRAS^G12C^ resulted in a significant decrease in cell viability (Figure 5F, 5G and 5H). Induction of KRAS^G12C^ also resulted in increased pERK after 24 hours in the three cell lines. The loss of viability resulting from increased KRAS^G12C^ was rescued by treating the cells with 1 nM trametinib or 1 nM AMG 510 (Figure 5I-K). Treatment with 10 nM or 1 nM AMG 510 and trametinib resulted in decreased pERK levels suggesting rescue may be due to buffering of ERK activity (Supplemental Figure 5E). This confirms that amplification of KRAS^G12C^ signaling can result in lethality in the absence of MEK inhibition, further implicating this as a determinant of trametinib addiction in our model system. These findings also suggest that KRAS signaling – and subsequently ERK activity – must be finely tuned for optimal lung cancer cell growth. Complete suppression of KRAS with siRNA or high concentrations of AMG 510 results in cell death (Supplemental Figure 5F-H). However, increased KRAS signaling through amplification of KRAS^G12C^ also leads to cell death through ERK hyperactivation, which can be rescued through buffering p-ERK to tolerable levels with modest concentrations of AMG 510 or trametinib. A similar phenomenon is observed in H358 sg*RB1*#4^tramR^. Cells are initially addicted to MAPK pathway signaling, and highly sensitive to MEK inhibitor treatment. In response to chronic treatment with trametinib, mutant KRAS becomes amplified and reactivates pERK signaling. When trametinib is removed, however, high levels of mutant KRAS signaling lead to excessive pERK and apoptosis (Figure 6). This balance of KRAS signaling and pERK levels leads to therapeutic vulnerabilities, which can be exploited to both prevent and counteract acquired MEK inhibitor resistance.

**Figure 6.**
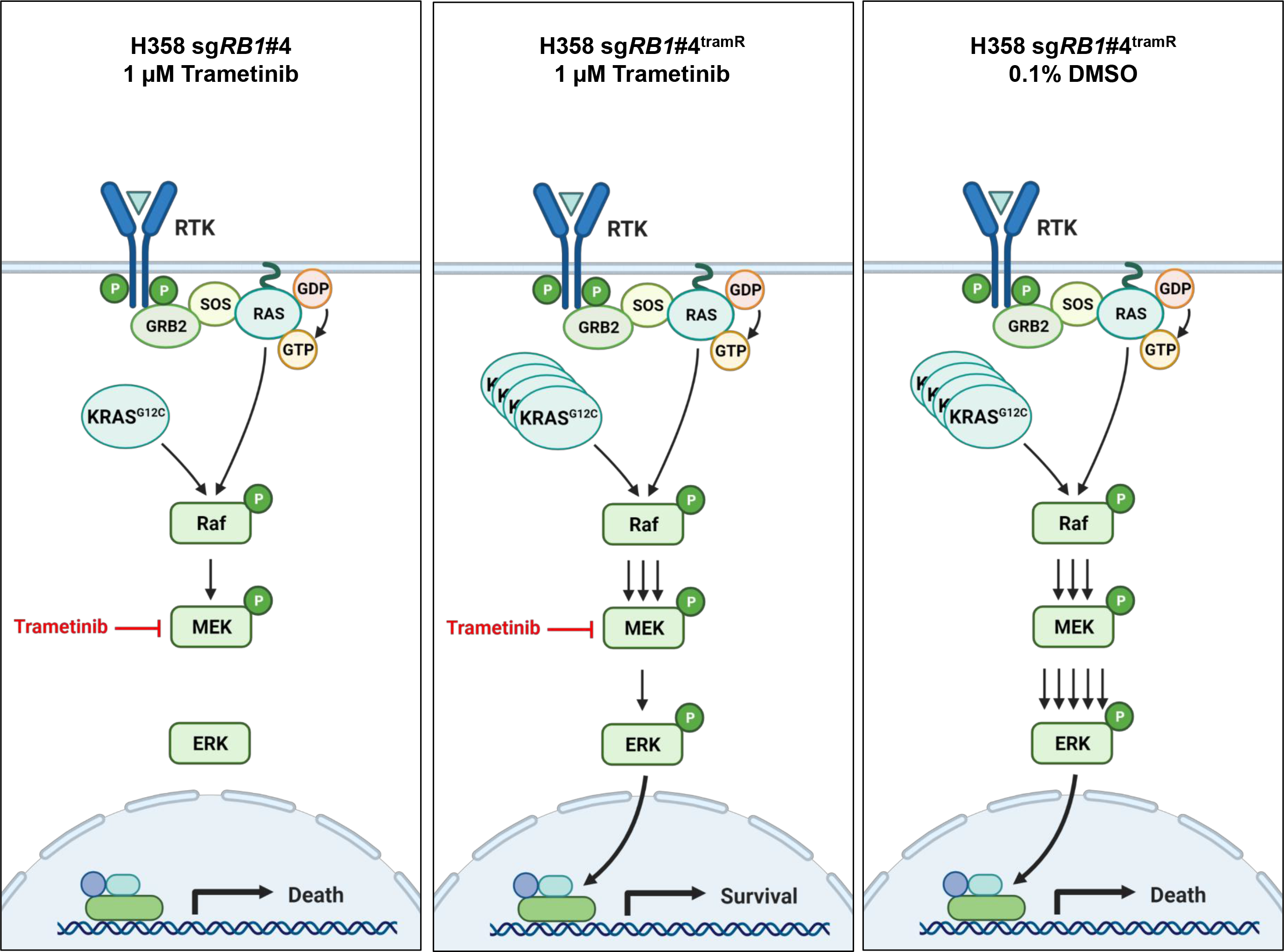
Mutant KRAS amplification is associated with resistance and dependence to trametinib. Parental H358 cells are sensitive to trametinib. In H358 sg*RB1*#4^tramR^ cells, KRAS^G12C^ amp is associated with resistance to trametinib. In these same cells, when trametinib is removed, KRAS^G12C^ amp drives ERK hyperactivation and cell death. Figure made with BioRender, adapted from “RAS Pathway”, by BioRender.com (2021). Retrieved from https://app.biorender.com/biorender-templates

## Discussion

Due to promising pre-clinical data, targeting the MAPK signaling pathway through MEK inhibition remains an attractive option for treatment of KRAS mutant LUAD, despite recent clinical setbacks (17, 18). Here we sought to model acquired resistance to MEK inhibitors in KRAS mutant LUAD cells through dose escalation in order to define strategies to increase treatment effectiveness. We observed upregulation of ERBB3 and FGFR1 (Supplemental Figure 1J), as well as increased AKT levels, suggesting cells employed previously described avenues of intrinsic resistance to bypass MEK inhibition (19,23,24,26). Increased expression of EMT genes, as was observed in some of our cells (Supplemental Figure 1J), has been associated with a more invasive phenotype (53) and could be studied further in our models of trametinib resistance. In H1792 sgControl^tramR^ cells, we noted increased pERK levels coincident with a *MAP2K1* F53V (Supplemental Figure 4B) missense mutation. *MAP2K1* F53 mutations have been previously documented in cancer patients and validated as functional driver mutations (54, 55). Our cell line bearing the *MAP2K1* F53V mutation may therefore provide important insight into the role of *MAP2K1* in driving MEK inhibitor resistance upon future investigation.

Based on previous observations, we aimed to assess the role of RB inactivation in the development of MEK inhibitor resistance and found that one H1792 RB knockout clone and RB proficient cells were equally sensitive to trametinib while the other H1792 *RB1* KO and both H358 *RB1* KO cells had modestly higher IC50s relative to the control line (Figure 1A, 1B, Table 1). This mirrors previous results linking *RB1* loss to trametinib resistance (28, 29), although we observed a lesser effect using CRISPR/Cas9 to knockout *RB1* instead of acute siRNA mediated knockdown as previously reported (28, 29). However, H358 *RB1* knockout and control clones both remained sensitive to low doses of trametinib with IC50s in the nanomolar range. In addition, H358 and H1792 clones developed resistance to trametinib at the same rate (Supplemental Figure 1B), regardless of RB status, and both control and RB knockout resistant clones were resistant to >10 µM trametinib (Table 1). This contrasts previous reports where RB deficient KRAS mutant H358 cells developed resistance to MEK inhibition faster than cells with normal RB levels (29). We observed that RB inactivation may slightly decrease trametinib sensitivity of parental H358 cells, but did not have an impact on acquired resistance to trametinib in our model system, which we confirmed by re-expressing RB in knockout cells with no observed effects on trametinib sensitivity (Supplemental Figure 3D and 3E).

Of greatest interest, one of the cell lines, H358 sg*RB1*#4^tramR^, was found to be both resistant to, and dependent on, trametinib for survival (Figure 2). In this cell line, trametinib removal resulted in induction of ER stress signaling and apoptosis (Supplemental Figure 3B). Further investigation revealed that continued suppression of pERK2 is required for survival of this cell line and that cell death following drug removal could be rescued by genetic or pharmacological inhibition of ERK2 (Figure 3B-D). We subsequently found that hyperactivation of ERK2 upon drug withdrawal was driven by amplification of the KRAS^G12C^ allele in this context (Figure 5D). We validated this by ectopic expression of KRAS^G12C^ in H358, H23 and H1792 cells, which inhibited cell viability in all instances (Figure 5F-K). Our observations add to a growing body of evidence demonstrating that hyperactive MAPK signaling, specifically through ERK2, is toxic to cancer cells, in particular those already dependent on this pathway for survival (41,42,46,56,57). The distinction between ERK1 and ERK2 signaling is clear in our model, as inhibition of ERK1 alone further decreases viability of H358 sg*RB1*#4^tramR^ cells when trametinib is removed, whereas ERK2 inhibition rescues this effect. Comparison of downstream targets of ERK1 vs ERK2 might provide insight into which effectors drive cell death upon hyperactivation and which pathways the cells are dependent on for growth and survival. RB status was not found to affect the drug addiction phenotype (Supplemental Figure 3D and 3E), suggesting that the genetic alterations resulting in drug addiction could also arise in cells without RB loss.

Our observations of “drug addiction” closely mirror reports from melanoma (39–43,58), where amplification of components of the MAPK pathway lead to BRAF and MEK inhibitor resistance and also results in dependence on continued ERK suppression for survival. Here we present the first instance of drug addiction resulting from *KRAS* amplification. Our previous work has established that oncogenic mutations in *EGFR* and *KRAS* are mutually exclusive in LUAD due to toxicity induced by excessive ERK signaling when co-expressed (46). Here, we build on this finding by demonstrating that genetic alteration otherwise toxic to cancer cells can develop *de novo* as a response to treatment with a MAPK pathway inhibitor. While these acquired genetic alterations, in our instance amplification of the heterozygous mutant KRAS^G12C^ allele, confer drug resistance, this is only possible due to continued MEK/ERK suppression by trametinib, and upon removal of the drug, these alterations result in lethality due to ERK hyperactivation. The observation of addiction to MEK inhibitors *in vitro* suggests that this phenotype may also develop in patients undergoing treatment with inhibitors of this pathway. In a melanoma xenograft model, resistance to vemurafenib, a BRAF^V600E^ specific inhibitor, was forestalled by using an intermittent dosing strategy (59). A similar approach of intermittent dosing of trametinib in patients known to have tumors with mutant *KRAS* amplification may also prolong drug response by both killing cells dependent on MEK for survival when on drug, and extinguishing drug resistant clones with toxic acquired genetic alterations. Probing for KRAS amplification may be an indicator of potential response to such “drug holiday” management.

Successful implementation of such a strategy will require further preclinical work and characterization of biomarkers indicative of hyperactivation, as well as careful planning of dosing timing in patients, to be successful. In melanoma, there are reports of tumors that initially acquired resistance to BRAF inhibitor, responding to a rechallenge following a period where therapy was discontinued, suggesting that this phenotype can arise in patients (60). As *KRAS* amplification resulting in drug addiction is an acquired mechanism of resistance, treatments schedules with longer intervals will likely be more effective. A recent phase 2 trial in melanoma compared continuous versus intermittent dosing of BRAF and MEK inhibitors and found intermittent dosing of inhibitors did not improve progression free survival (61). In preclinical models of melanoma, drug addiction occurred only after *BRAF*^V600E^ was amplified to a level where it activated the MAPK pathway beyond toleration. In the above trial, the investigators did not assess *BRAF* amplification status in patients before removing them from drug and follow a dosing schedule standardized for imaging. Different patients may develop drug-addicted cells at different rates and removing drug for patients without sufficient *BRAF* amplification to promote drug addiction may instead promote tumor growth. Personalized timing based on assessment of mutant *BRAF* or *KRAS* amplification levels by sampling cfDNA for example, may be required to better elicit MAPK hyperactivation to forestall drug resistance using an intermittent dosing strategy in lung and melanoma patients. Additionally, evaluation of the frequency of mutant *BRAF* or *KRAS* in subsequent biopsies should also be used to inform a decision to halt treatment. If there is only a small subpopulation of the tumor that is drug addicted, there will be only minor effects after drug withdrawal.

Our discovery of drug addiction as a result of MEK inhibition has implications for both the treatment of KRAS mutant lung cancers and the continued study of MAPK pathway activation as a potential therapeutic target. Although MEK inhibitors alone or in combination with standard chemotherapy have not proven effective in the clinic, these compounds are still being investigated in combination with other targeted agents. With the FDA recently approving the first KRAS^G12C^ specific inhibitor, instances of clinical resistance are already being reported (62, 63). Importantly, the reactivation of RAS-MAPK signaling has been reported as a key mechanism by which tumors overcome KRAS^G12C^ inhibition in this context. For this reason, trametinib is currently being tested in combination with AMG 510 in clinical trials (NCT04185883) in order to block reactivation of MAPK signaling and the resulting resistance. Combination of KRAS specific inhibitors with MEK inhibitors may sensitize cells that initially displayed intrinsic resistance to MEK inhibitors alone, which would result in more cases of adaptive resistance to MAPK pathway inhibition. This is analogous to the use of MEK inhibitors with BRAF targeted therapy in melanoma, a setting where drug addiction has been reported (40), underscoring the continued importance of defining avenues of trametinib resistance in LUAD. Probing for *KRAS* amplification in patients treated with MEK inhibitors alone or in combination with other therapies may help identify those who might benefit most from a drug holiday. Our cell line provides a model system for further study into how drug addiction may develop in patients, as well as how we can induce or further potentiate the effects of hyperactive ERK2 by inhibiting negative regulators of that pathway, such as DUSP6 (46).

## Author contributions

Conceptualization, D.A.F., Y.I., A.M.U. and W.W.L.; Methodology, D.A.F., Y.I., F.D.J., A.M.U, R.S., M.L. and W.W.L.; Investigation, D.A.F., Y.I., F.D.J., G.D.R., R.S., D.L., R.S., M.L., and W.W.L.; Formal Analysis, D.A.F., Y.I., and W.W.L ; Visualization, Y.I., A.N., D.A.F., and W.W.L.; Writing – Original Draft, D.A.F. and W.W.L.; Writing – Review & Editing, D.A.F., Y.I., R.S., M.L., A.M.U. and W.W.L.; Funding Acquisition, W.W.L.; Resources, M.L. and W.W.L; Supervision, W.W.L.

